# Degraded mapping of disparity tuning in visual cortex explains deficits in binocular depth perception

**DOI:** 10.1101/2025.04.22.649872

**Authors:** Thomas C Brown, Carrie Barr, Mason Antin, Vy Nguyen, Aaron W McGee, Jason M Samonds

**Affiliations:** Translational Neurosciences, University of Arizona College of Medicine - Phoenix; Department of Neuroscience, University of Texas at Austin; Center for Learning & Memory, University of Texas at Austin

## Abstract

Sensory cortex is highly organized, but the function of this organization is contentious. We show that closing one eye during the developmental critical period in juvenile mice (monocular deprivation) caused lasting impairments in binocular depth perception and disrupted cortical maps of binocular disparity tuning. In normal mice, disparity tuning was concentrated in the central visual field, particularly with respect to azimuth and in higher visual areas (HVAs). Additionally, neurons transitioned from encoding nearer to farther stereoscopic depths from the center toward the peripheral visual field, especially along elevation and within HVAs. Monocular deprivation did not reduce overall disparity selectivity across the neuronal population, but rather weakened the retinotopic and hierarchical organization of disparity tuning and reduced trial-to-trial reliability of neuronal responses. Analysis of single-trial responses from neurons tuned to near disparities showed that this reduced response reliability is sufficient to explain the impaired depth discrimination observed in deprived mice.

## Introduction

Hubel and Wiesel demonstrated that the selectivity of neurons in the primary visual cortex (V1) of cats and non-human primates is organized with respect to retinotopy, orientation, and ocular dominance^1,2^. Several decades of subsequent research have identified the spatial organization or clustering of functional characteristics of neurons in numerous brain regions of many mammals^3,4^. Yet, not all ‘tuning’ properties of cortical neurons are organized in this manner in all mammals. Crucially, whether the mapping of tuning properties serves a functional purpose is controversial^5-7^.

Neurons in the region of binocular overlap in V1 are tuned for ocular dominance, the preference for visual input to one of the eyes. Ocular dominance is disrupted by closing one eye (monocular deprivation, MD) during a critical period of visual development. In juvenile mammals, including non-human primates, cats, and rodents, MD shifts responsiveness of binocular neurons in V1 to favor the open eye and changes the map of ocular dominance^8-12^. MD during the critical period has been widely used as a model of amblyopia, a visual disorder that often presents with deficits in acuity and binocular depth perception^13-16^. In the mouse, MD for only 4-8 days causes maximal shifts in the ocular dominance of neurons in V1^11,12^. However, re-opening the deprived eye results in near complete recovery of normal ocular dominance within a few days^17^. In contrast, MD lasting 30 days that spans the critical period causes enduring deficits in both tuning for ocular dominance as well as visual acuity that persist after binocular vision is restored^18^.

Neurons in the region of binocular overlap in V1 may also be tuned for disparity, the difference between the position of an image on the retina arising from the horizontal offset of the two eyes^19^. Tuning for ocular dominance does not correlate with tuning for disparity in non-human primates, cats, or mice^20-22^. Tuning for disparity is more relevant than tuning for ocular dominance for binocular depth perception^19^. In contrast to cats and non-human primates, neurons with similar orientation and disparity tuning are generally not clustered in V1 of the mouse^23-25^. However, the average disparity preference in V1 and higher visual areas (HVAs) varies systematically across elevation representing on average nearer to farther depths from the lower to upper visual field, respectively^26^. This provides a map in the mouse based on disparity to test for functional significance that is analogous to a map found in V1 of non-human primates^27^.

One study has measured disparity selectivity in V1 of mice following 8 days of MD and found weaker selectivity, but that was measured in anesthetized mice immediately after the deprived eye was opened^28^, so it is unknown if this deficit would have resolved upon restoring binocular vision or how the deficit relates to binocular depth perception. Recently, we developed a quantitative behavioral assay for binocular depth perception^29^. Here, we examined if 30 days of MD in juvenile mice followed by several weeks of binocular vision caused enduring impairments in binocular depth discrimination and how it perturbed the representation and organization of disparity tuning in V1 and HVAs. Comparing these findings revealed that deficits in binocular depth perception were not associated with reduced disparity selectivity, but were rather associated with degraded mapping and reduced reliability of responses by disparity-tuned neurons in V1 and especially HVAs.

## Results

Mice depend on binocular vision to preferentially exit to the nearest platform when the difference in distances is as small as 13 cm and as large as 58 cm in the pole descent cliff task (PDCT) (Figure 1a)^29^. We tested if MD spanning the critical period of ocular dominance plasticity and maturation of visual acuity impairs performance in this task^12,18^. Mice received MD by suturing closed one eye at post-natal (P) day 22 for 30 days. The suture was then removed and binocular vision restored for 6 or more weeks^18,30^. Mice receiving 30 days of MD displayed significantly worse performance than mice with normal binocular development (non-deprived) when the near platform was positioned 2.5 cm below the choice point and the three other quadrants were 15.2 cm away (Figure 1b, left). The magnitude of the impairment in performance was smaller, but still significant, for mice tested with larger differences in depth between the platforms (Figure 1b, right, 2.5 cm versus 30.5 cm). Mice receiving 30 days of MD did not take longer to descend the pole compared to non-deprived mice suggesting that the deficit was due to discrimination of depth and not general to sensorimotor behavior (Extended Data Figure 1). Enduring deficits from MD were not observed when mice were tested using the classical visual cliff task (Extended Data Figure 2). This is likely because the PDCT places surfaces in a part of the visual field with a greater proportion of binocular overlap making it superior for detecting binocular depth discrimination deficits^29^.

**Figure 1.**
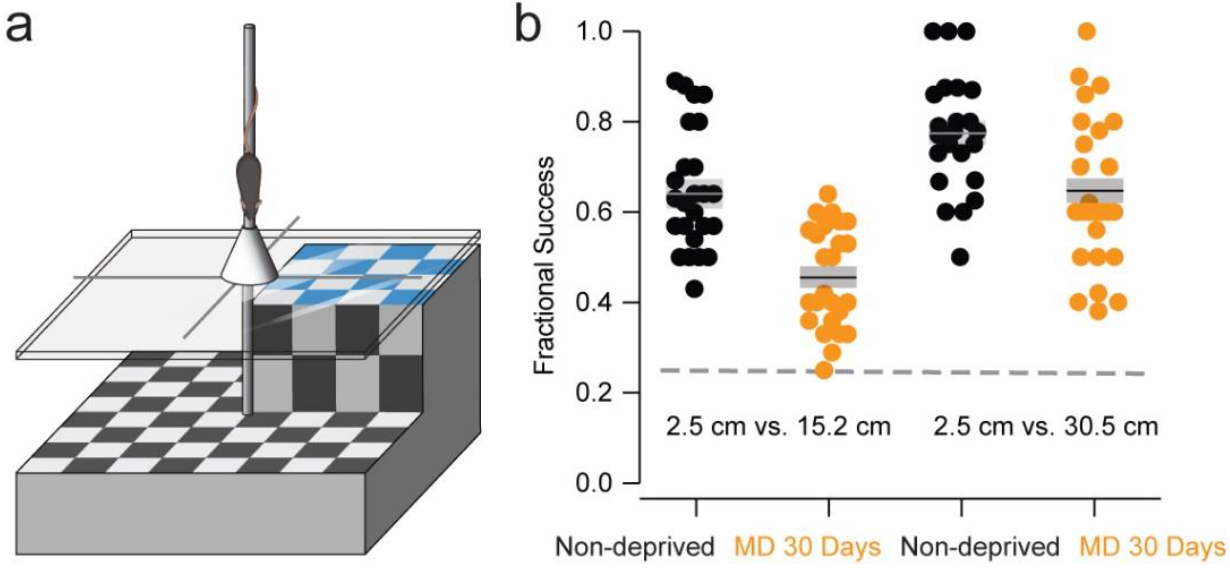
Long-term monocular deprivation impaired binocular depth discrimination. **a**, A cartoon of the pole descent cliff task used to measure binocular depth discrimination in non-deprived and 30-day MD mice. **b**, The Fractional Success is the fraction of trials that each mouse chose the near surface (2.5 cm, blue in **a**) when compared to surfaces 15.2 and 30.5 cm away. This fraction was significantly reduced by MD for platforms set at distances of 2.5 cm vs. 15.2 cm (Non-deprived, n = 25; MD 30 days, n=25; bootstrap test, p < 0.0001) and 2.5 cm vs. 30.5 cm (Non-deprived, n = 23; MD 30 days, n = 26; bootstrap test, p = 0.005). Horizontal lines for each group represent the mean and shading is standard error of the mean. Circles represent individual mice. The dashed horizontal line indicates performance at chance levels of Fractional Success (0.25).

To understand why mice receiving 30 days of MD had a deficit in binocular depth discrimination, we examined responses in excitatory cortical neurons in mice that expressed the genetically-encoded calcium indicator GCaMP6s^31^. We employed two-photon microscopy to measure the response of excitatory neurons in L2/3 to a range of stereoscopic depths. Depth was produced by varying the binocular disparity (-9.2 to +9.2°) of dynamic random dots projected within a central region that covered the vertical extent of the screen (100° of visual angle) and the entire horizontal width of binocular overlap in the mouse (45.5° of visual angle)^32^. This central vertical strip was flanked on both sides by dynamic random dots with zero disparity. The visual stimulus produced stereoscopic surfaces in front of and behind the position of the screen (Figure 2a). The range of stereoscopic depths presented was greater than the range of depths mice encountered when performing the PDCT^29^. We imaged neurons in the primary visual cortex (V1), as well as the HVAs of the lateromedial (LM) and rostrolateral (RL) visual cortex that represent the upper and lower visual field, respectively.

**Figure 2.**
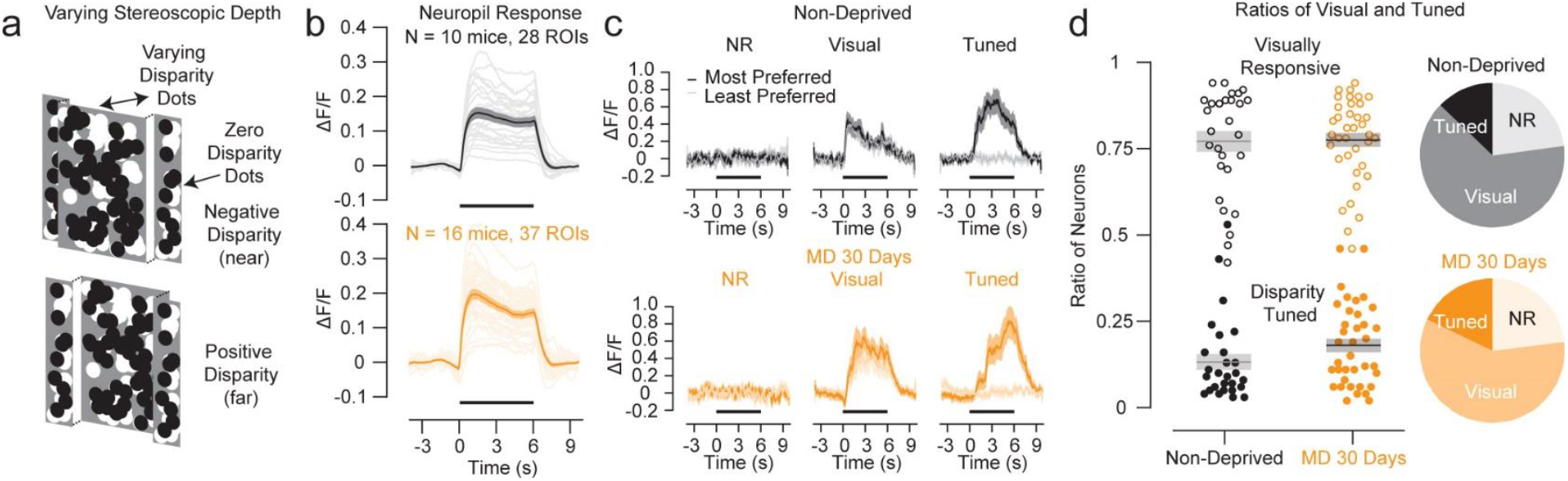
Responses to stereoscopic stimuli were similar between non-deprived mice and 30-day MD mice. **a**, A cartoon of the stereoscopically rendered surfaces shown to the mice. **b**, Neuropil responses for regions-of-interest (ROIs) (light lines) and the average (dark line) were similar between non-deprived and 30-day MD mice. **c**, All identified neurons in two-photon microscopy were classified into three groups: non-responsive (NR, no significant visual response), visual (significant visual response, sign test: stimulus > pre-stimulus response, p < 0.05), and tuned (significant disparity tuning, Kruskal-Wallis test, p < 0.05). Examples of the temporal response (n = 10 repeats for each disparity) for each neuron type were plotted for two mice for the disparities that resulted in the largest (most preferred) and smallest (least preferred) responses. The black horizontal bar represents when the stimulus was displayed. **d**, Ratios of significant visually-responsive (open circles, visual plus tuned) and significant disparity-tuned (solid circles, tuned) neurons were similar between non-deprived and 30-day MD mice (bootstrap test, p = 0.91 and p = 0.11, respectively). All shaded regions are standard error of the mean.

First, we compared responsiveness between non-deprived mice and mice that experienced 30 days of MD. There were not obvious differences in visually-evoked and disparity-selective responses between non-deprived mice and 30-day MD mice. The neuropil response (see Materials and Methods) represents the aggregate of neuronal activity within a region of interest (ROI) and there was no evidence of a reduced responsiveness to stereograms for 30-day MD mice compared to non-deprived mice (Figure 2b). We classified individual neurons as (1) non-responsive to our stereoscopic stimuli (Figure 2c, “NR”), (2) significantly visually-responsive but not selective for disparity (Figure 2c, “Visual”), and (3) significantly disparity-tuned (Figure 2c, “Tuned”). The ratios of visually-responsive (visual plus tuned) or disparity-tuned neurons were also not significantly different between non-deprived compared to 30-day MD mice (Figure 2d). Overall, these results suggest that in the least, 30-day MD mice should potentially have sufficient visual responses and disparity selectivity to support binocular depth discrimination comparable to non-deprived mice.

Although we always targeted neurons well within the region of binocular overlap, that cortical region is large enough and our stimulus was large enough to allow us to sample across a wide range of retinotopic locations and multiple cortical visual areas. This allowed us to test for differences between non-deprived and 30-day MD mice specific to retinotopy and area. To examine the retinotopic and cortical organization of the ratio of visually-responsive and disparity-tuned neurons, we aligned imaging windows with respect to the V1/RL/LM border for each mouse identified by retinotopic mapping with optical imaging of calcium responses^33^ (Figure 3a, red dot). We then measured the distance perpendicular from the RL/LM border (negative: lower V1 or RL, positive: upper V1 or LM) and the distance perpendicular from the V1/HVA border (negative: V1, positive: RL, LM, AL) for each neuron. We generated a complete two-dimensional cortical map of the ratio of significant visually-responsive (Figure 3b) and disparity-tuned (Figure 3e) neurons within sliding ±0.25 mm windows every 0.01 mm along these axes. Only pixels where the window included at least 100 neurons in both non-deprived and 30-day MD mice were included in each map. For reference, we overlaid these maps with the average location of visual area borders (solid white) and retinotopic contours (dashed white) aligned to our reference point^33^. To quantify and assess the statistical significance of any differences of maps between non-deprived and 30-day MD mice, we computed the ratio of significantly visually-responsive neurons and the ratio of significant disparity-tuned neurons for each ROI with respect to either elevation or azimuth. Elevation or azimuth was determined for each neuron by translating their cortical location into retinotopic coordinates using an average retinotopic map of visual cortex^33^. Ratios were then computed in windows of 0° to ≤2.5°, >2.5° to ≤5°, >5° to ≤10°, and >10° to ≤15° for elevation or azimuth and separately for V1 and HVAs. We then computed the average ratio of visually responsive or disparity-tuned neurons across ROIs in each case.

**Figure 3.**
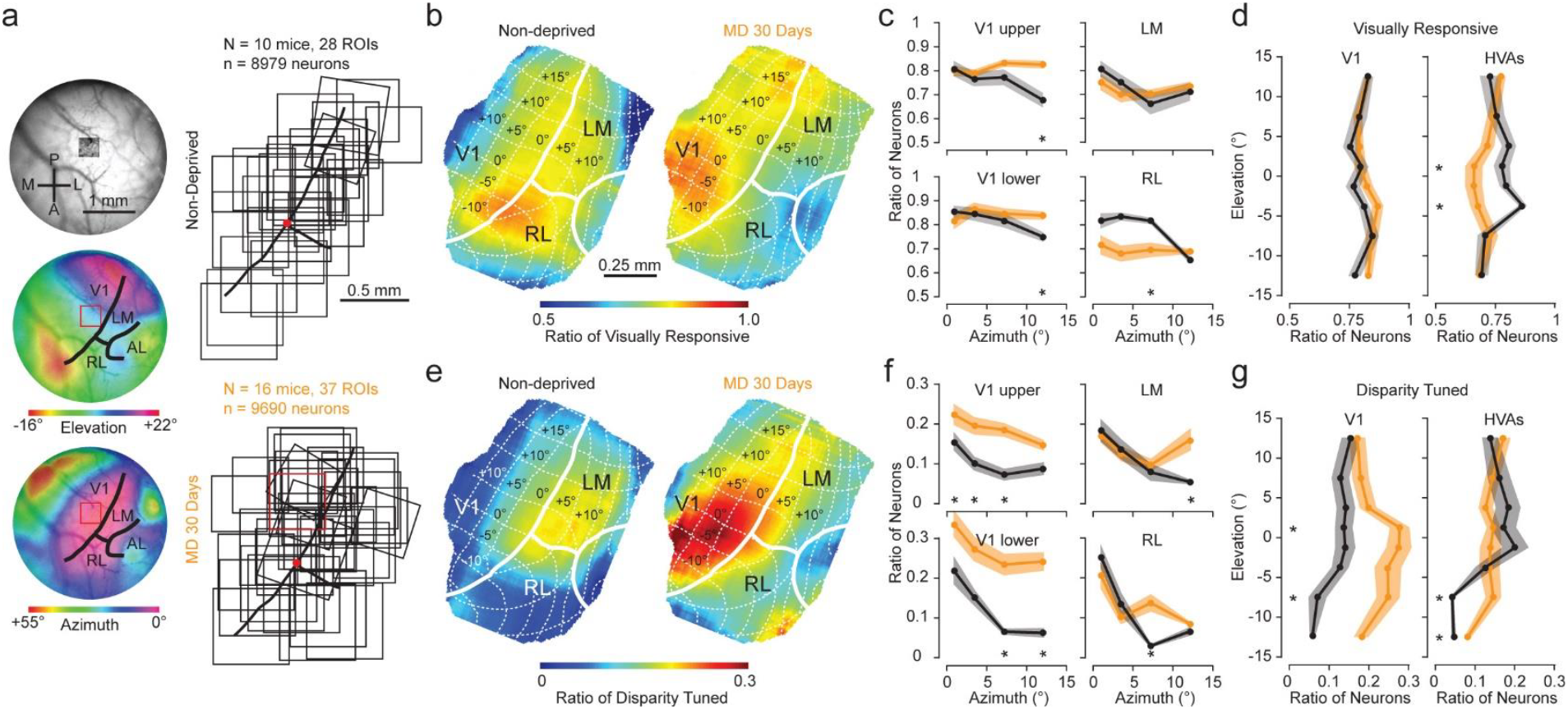
Monocular deprivation altered the spatial representation of visual responsiveness and disparity tuning in binocular visual cortex. **a**, Left, for each mouse, we aligned two-photon (2P) microscopy regions of interest (ROI) with the overall vasculature. Then, we mapped retinotopy to determine the binocular region borders of V1, RL, and LM to align ROI to a single point. Right, all borders and ROIs (400-550 µm) for either non-deprived or 30-day MD mice were aggregated and aligned within a single map at this point (N = 28 and 37, respectively). Anterior-Posterior and Lateral-Medial axes indicated in top left image. **b**, The color of each pixel in the map represents the ratio of significant visually-responsive neurons within a ±0.25 mm window for all ROIs for non-deprived mice combined. Solid white lines represent the area borders and dashed white lines represent retinotopic contours based on average locations for mice^33^. **c**, Ratio of significant visually-responsive neurons (weighted mean for all ROIs based on number of neurons) for non-deprived mice (black) and 30-day MD mice (orange) with respect to azimuth in upper V1 (upper left, bootstrap ANOVA, p = 0.04 and 0.23, for non-deprived and 30-day MD mice, respectively), lower V1 (lower left, p = 0.03 and 0.50, respectively), LM (upper right, p = 0.21 and 0.40, respectively), and RL (lower right, p < 0.0001 and p = 0.76, respectively). **d**, Ratio of significant visually-responsive neurons for non-deprived mice (black) and 30-day MD mice (orange) with respect to elevation in V1 (left, bootstrap ANOVA, p = 0.23 and 0.17, respectively) and HVAs (right, p = 0.0003 and 0.11, respectively). **e**, Same maps as **b**, but pixel color represents the ratio of significant disparity-tuned neurons. **f**, Ratio of significant disparity-tuned neurons for non-deprived mice (black) and 30-day MD mice (orange) with respect to azimuth in upper V1 (upper left, bootstrap ANOVA, p = 0.02 and 0.03, respectively), lower V1 (lower left, p = 0.002 and 0.31, respectively), LM (upper right, p < 0.0001 and p = 0.08, respectively), and RL (lower right, p < 0.0001 and p = 0.02, respectively). **g**, Ratio of significant disparity-tuned neurons for non-deprived mice (black) and 30-day MD mice (orange) with respect to elevation in V1 (left, bootstrap ANOVA, p < 0.0001 and p = 0.07, respectively) and HVAs (right, p < 0.0001 and p = 0.24, respectively). Significant differences (bootstrap test, p < 0.05) between non-deprived and 30-day MD data points are noted with an asterisk. Shaded regions are standard error of the mean.

For non-deprived mice, the ratio of visually-responsive neurons was highest at the V1/RL/LM borders that represent the center of the visual field and dropped rapidly to 0.5 with respect to azimuth (Figure 3b, left). This falloff was statistically significant in binocular visual cortex, except for LM (Figure 3c, black). In contrast, retinotopic preference for higher ratios of visually-responsive neurons was broader in 30-day MD animals and more prominent in V1 compared to HVAs (Figure 3b, right). The ratio of visually-responsive neurons did not significantly depend on azimuth for 30-day MD mice in any area of binocular visual cortex (Figure 3c, orange). In V1 for mice with 30 days of MD, the ratio of visually-responsive neurons remained high across azimuth and was significantly greater compared to non-deprived mice for retinotopic locations farther from the center of the visual field (Figure 3c, left, asterisks). In HVAs for mice with 30 days of MD, the ratio of visually-responsive neurons remained low across azimuth and was significantly lower in RL for retinotopic locations closer to the center of the visual field (Figure 3c, right, asterisks). In V1, the ratio of visually-responsive neurons did not significantly depend on elevation for non-deprived or 30-day MD mice and there were no significant differences in the ratios of visually-responsive neurons between non-deprived and 30-day MD mice in V1 (Figure 3d, left). In HVAs, the ratio of visually-responsive neurons did significantly depend on elevation for non-deprived mice (Figure 3d; right, black), but not for 30-day MD mice (Figure 3d; right, orange). There were significantly higher ratios of visually-responsive neurons for non-deprived compared to 30-day MD mice for HVA neurons representing the lower central portion of the visual field (Figure 3d; right, asterisks). Overall, this reveals that 30 days of MD significantly altered the mapping of visually-responsive neurons in V1 and HVAs and demonstrates that visual responsiveness in HVAs in these mice was impaired.

For both non-deprived and 30-day MD mice, the highest ratio of disparity-tuned neurons were restricted in proximity to the V1/RL/LM borders that represent the central portion of the visual field (Figure 3e). For non-deprived mice, this enhanced ratio of disparity-tuned neurons dropped rapidly to near zero with eccentricity and appeared to be more prominent in the HVAs (LM and RL) compared to V1 (Figure 3e, left). Retinotopic preference for higher ratios of disparity-tuned neurons was broader in 30-day MD animals always remaining above 0.1 and was more prominent in V1 compared to HVAs (Figure 3e, right). The rapid fall-off in the ratio of disparity-tuned neurons was very evident and statistically significant with respect to azimuth for non-deprived mice everywhere in binocular visual cortex (Figure 3f, black). A higher ratio of disparity-tuned neurons for 30-day mice was more spread out, but still significantly depended on azimuth in some cases (Figure 3f, orange). The ratio of disparity-tuned neurons was significantly higher for 30-day MD mice compared to non-deprived mice throughout V1 (Figure 3f, left, asterisks). These ratios were generally lower for 30-day MD mice in the HVAs and only significantly higher than non-deprived mice for retinotopic locations farther from the center of the visual field (Figure 3f, right, asterisks). The dependence of the ratio of disparity-tuned neurons on eccentricity was also much clearer with respect to elevation for non-deprived mice compared to mice with 30 days of MD. In V1 and HVAs, the ratio of disparity-tuned neurons did significantly depend on elevation for non-deprived mice (Figure 3g; black), but did not for 30-day MD mice (Figure 3g, orange). This was especially obvious in the HVAs where the highest ratio of disparity-tuned neurons was in the lower central portion of the visual field (Figure 3g, right, black). There was a significantly higher ratio of disparity-tuned neurons for mice with 30 days of MD compared to non-deprived mice throughout V1, but only for neurons representing the lowest portions of the visual field in HVAs (Figure 3g, asterisks). Overall, this supports that 30 days of MD significantly altered the mapping of disparity-tuned neurons in V1 and HVAs with a more severe difference in HVAs.

Although 30 days of MD clearly changed the representation of disparity in mice, it was not clear that anything we described thus far would result in reduced disparity discrimination. If anything, 30-day MD mice had a greater representation of disparity-tuned neurons compared to non-deprived mice, especially in V1. We hypothesized that impaired disparity discrimination in 30-day MD mice could be caused by an underrepresentation of neurons tuned for disparities relevant to the depths discriminated by mice in the PDCT. Therefore, we examined whether 30 days of MD changed which disparities were represented in the visual cortex. We rejected this hypothesis because we determined that 30-day MD mice had neurons tuned for disparities that covered the entire range of disparities represented in non-deprived mice (Figure 4a) and there was no significant difference between their distributions of preferred disparities (Figure 4b). Therefore, we tested for differences in the representation of preferred disparity specific to retinotopy and cortical area.

**Figure 4.**
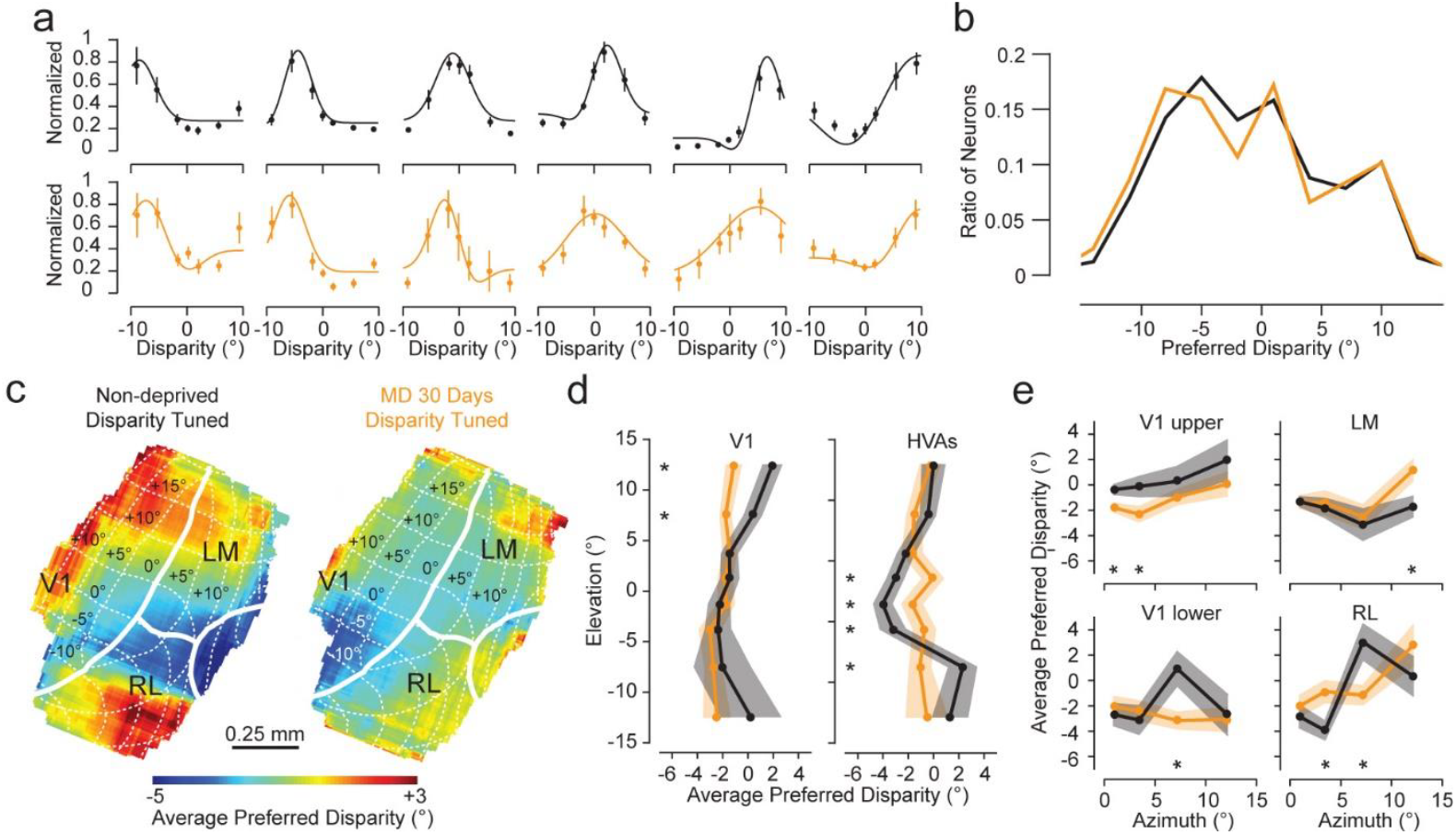
Monocular deprivation severely degraded mapping of preferred disparity in binocular visual cortex. **a**, Both non-deprived (top row) and 30-day MD (bottom row) mice had preferred disparities that covered the entire range of disparities that we tested (±9.2°). **b**, Distributions of preferred disparities were not significantly different between non-deprived (black) and 30-day MD mice (orange) (mean ± SD, -1.24 ± 6.71° and -1.58 ± 7.03°; n = 1152 and 1737, respectively; bootstrap test, p = 0.20). **c**, Each point in the map represents the average preferred disparity of significant disparity-tuned neurons within a ±0.25 mm window for either non-deprived (left) or MD (right) mice. Each pixel in the maps included at least 10 significant disparity-tuned neurons in both non-deprived and 30-day MD mice. **d**, Mean preferred disparity for non-deprived mice (black) and 30-day MD mice (orange) with respect to elevation for neurons in V1 (left, bootstrap ANOVA, p = 0.0002 and 0.17, respectively) and HVAs (right, p < 0.0001 and p = 0.21, respectively). **e**, Mean preferred disparity for non-deprived mice (black) and 30-day MD mice (orange) with respect to azimuth for neurons in upper V1 (upper left, one-way ANOVA, p = 0.14 and 0.08, respectively), lower V1 (lower left, p = 0.01 and 0.46, respectively), LM (upper right, p = 0.17 and 0.002, respectively), and RL (lower right, p < 0.0001 and p = 0.01, respectively). Significant differences (bootstrap test, p < 0.05) between non-deprived and 30-day MD data points are noted with an asterisk. Error bars and shaded regions are standard error of the mean.

Based on a previous study in mice^26^, the average preferred disparity depends on elevation within binocular cortical areas, so we tested if MD caused changes to that map. In both non-human primates and mice, the distribution of preferred disparity for neurons transitions from representing closer depths to farther depths for receptive fields along the lower to upper visual field^26,27^. We measured similar mapping for the average preferred disparity (Figure 4c, left). The map correlated with the map of the ratio of disparity-tuned neurons. The central part of the visual field that has a greater ratio of disparity-tuned neurons that represent on average negative disparities or near stereoscopic surfaces (Figure 4c, left, blue). Disparity-tuned neurons surrounding this region represent on average more positive disparities or stereoscopic surfaces that are farther away (Figure 4c, left, red). Similar to the map of the ratio of disparity-tuned neurons, 30 days of MD appeared to result in a degraded map, especially in HVAs (Figure 4c, right). When we computed the average preferred disparity for neurons with respect to either elevation in V1 or HVAs, the average preferred disparity for non-deprived mice depended significantly on elevation (Figure 4d, black), but did not for 30-day MD mice (Figure 4d, orange). The map with respect to elevation appears to strengthen from V1 to HVAs for non-deprived mice (Figure 4d, black, left-to-right) and weaken from V1 to HVAs for 30-day MD mice (Figure 4d, orange, left-to-right). The map for average preferred disparity with respect to azimuth was not as clear in either non-deprived or 30-day MD mice. In general, farther disparities were represented in neurons that were more distant from the central part of the visual field (Figure 4e). Overall, this reveals that 30 days of MD significantly altered the mapping of preferred disparity in V1 and HVAs, yet how this would impair depth discrimination remained unclear.

Therefore, we examined how degrading the map of the ratio of disparity-tuned neurons and average preferred disparity could reduce disparity discrimination performance. Our first hypothesis was that mapping was important for enhancing disparity selectivity because previous studies have shown that altered visual experience during the critical period can both degrade neuronal orientation selectivity and the map of orientation preference^34^. Additionally, one previous study reported that mice receiving 8 days of MD had reduced disparity selectivity upon eye re-opening^28^. We measured if disparity-tuned neurons were more selective in non-deprived mice compared to 30-day MD mice. We measured the strength of disparity tuning for disparity-tuned neurons in response to random dot stereograms using the same vector-based disparity selectivity index employed in a previous study (DSI, see Materials and Methods)^28^. The DSI vector is 0 for neurons when the responses to all disparities are equal and 1 when there is a response to only a single disparity. Both non-deprived and 30-day MD mice had a wide range of DSI values (Figure 5a). For both non-deprived and 30-day MD mice, average DSI was similar and strongest in the upper central visual field, but the stronger DSI generally spread out over more distance in mice with 30 days of MD (Figure 5b, right versus left). When we computed the average DSI for neurons with respect to either elevation or azimuth, DSI depended significantly on retinotopy for both non-deprived and 30-day MD mice in both V1 and HVAs (Figure 5c-d). However, the mapping was relatively broad for average DSI overall. Generally, significant differences between non-deprived and 30-day MD mice were actually in the opposite direction of the prediction with stronger DSI observed in 30-day MD mice (Figure 5c-d, orange vs black, asterisks). Therefore, 30 days of MD did not cause any enduring deficits in the selectivity of disparity tuning for individual neurons.

**Figure 5.**
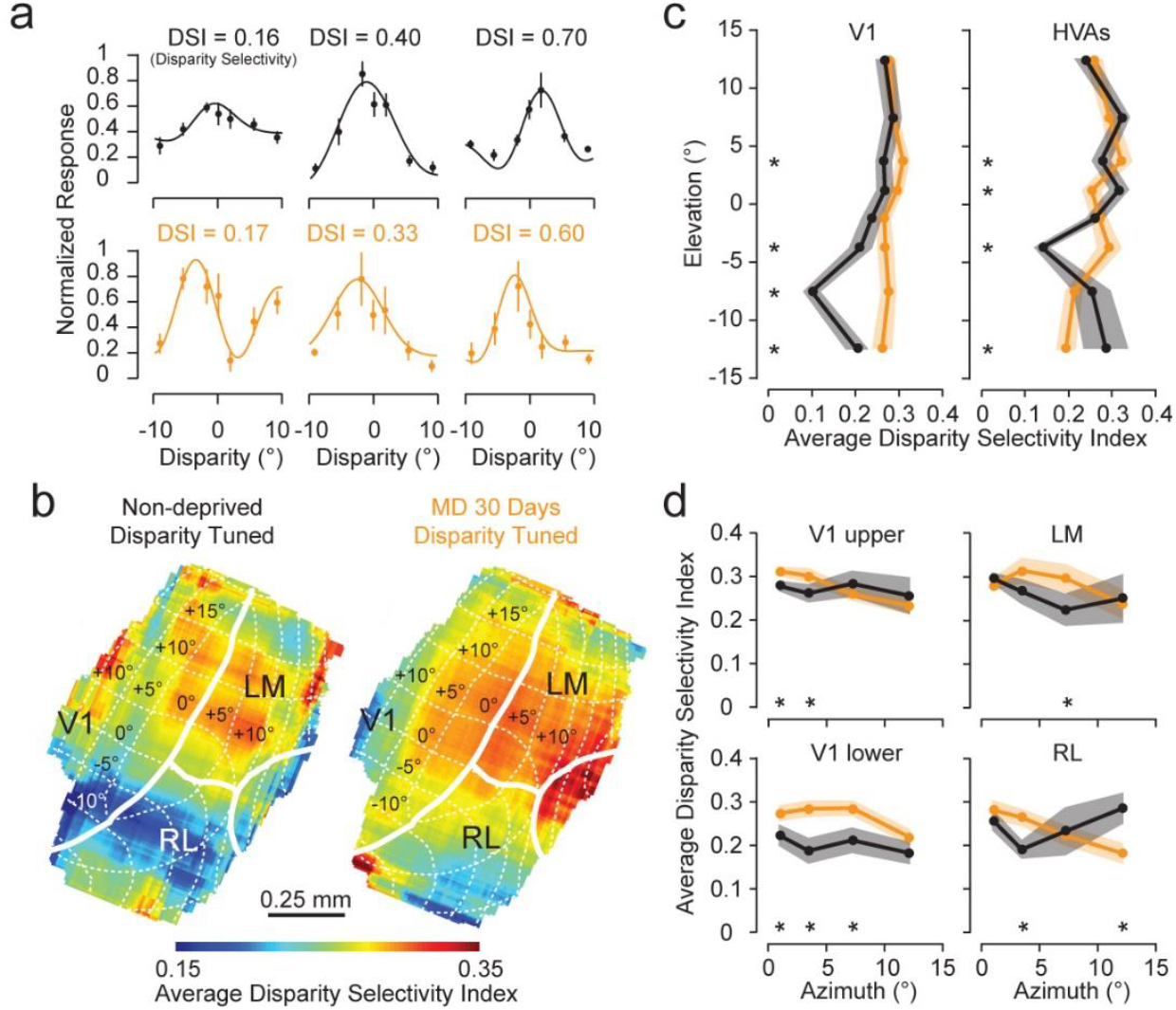
Monocular deprivation did not reduce disparity selectivity. **a**, The disparity selectivity index (DSI) represents how strongly the neuron responds to a single disparity relative to other disparities. The non-deprived (black) and 30-day MD (orange) mice had a similar range of weak to strongly disparity-tuned neurons **b**, Each point in the map represents the average disparity selectivity index (DSI) of significant disparity-tuned neurons within a ±0.25 mm window for either non-deprived (left) or MD (right) mice. Each pixel in the maps included at least 10 significant disparity-tuned neurons in both non-deprived and 30-day MD mice. **c**, Mean DSI for non-deprived mice (black) and 30-day MD mice (orange) with respect to elevation for neurons in V1 (left, bootstrap ANOVA, p < 0.0001 and p = 0.08, respectively) and HVAs (right, p < 0.0001 and p < 0.0001, respectively). **d**, Mean DSI for non-deprived mice (black) and 30-day MD mice (orange) with respect to azimuth for neurons in upper V1 (upper left, one-way ANOVA, p = 0.47 and 0.0003, respectively), lower V1 (lower left, p = 0.28 and 0.005, respectively), LM (upper right, p = 0.07 and 0.14, respectively), and RL (lower right, p = 0.06 and p < 0.0001, respectively). Significant differences (bootstrap test, p < 0.05) between non-deprived and 30-day MD data points are noted with an asterisk. Error bars and shaded regions are standard error of the mean.

An alternative hypothesis we had was that cortical organization with respect to retinotopy and hierarchy was important to generate reliable trial-to-trial disparity-dependent responses. One recent study found that reliability of trial-to-trial orientation-selective responses increased over the critical period of visual development^35^. We evaluated whether 30 days of MD reduced trial-to-trial reliability of disparity-dependent responses and if so, how that reduction related to the organization of disparity tuning that we observed in Figures 3 and 4.

To this point, we conducted all of our analyses on neuronal responses averaged over 10 repeats to the same disparities. However, natural stereoscopic depth discrimination involves a moment-by-moment estimate of disparity. Figure 6a illustrates lower trial-to-trial variation in disparity-dependent responses for an example neuron in a non-deprived animal (black) compared to an example neuron in a 30-day MD animal (orange). This difference is also clear in the larger error bars of disparity-dependent responses for several example neurons in Figures 4a and 5a for 30-day MD mice (orange) compared to non-deprived mice (black). We quantified the signal-to-noise ratio (SNR) for each neuron by taking the maximum mean response divided by the standard deviation. Unlike DSI, SNR was clearly stronger in non-deprived compared to 30-day MD mice (Figure 6b, left versus right). Additionally, cortical mapping of average SNR for non-deprived mice (Figure 6b, left) appeared to correlate with the map of the ratio of disparity-tuned neurons (Figure 3d) and average preferred disparity (Figure 4c). The highest SNR for non-deprived mice was observed in the center of the visual field and more prominently in the HVAs, which corresponds to higher ratios of disparity-tuned neurons and neurons that prefer near stereoscopic depths. This organization was degraded in mice following 30 days of MD (Figure 6b, right). SNR significantly depended on elevation for non-deprived mice in both V1 and HVAs (Figure 5c, black). SNR was significantly greater in non-deprived compared to 30-day MD mice in the lower central part of the visual field (Figure 6c, asterisks). The organization of SNR with respect to azimuth was less clear, but generally SNR was significantly greater for non-deprived compared to 30 day MD mice closer to the center of the visual field (Figure 6d, asterisks). Overall, SNR appeared to most closely correlate with the organization of average preferred disparity so that near disparities were most reliably represented in the lower central portion of the visual field for HVAs in non-deprived mice.

**Figure 6.**
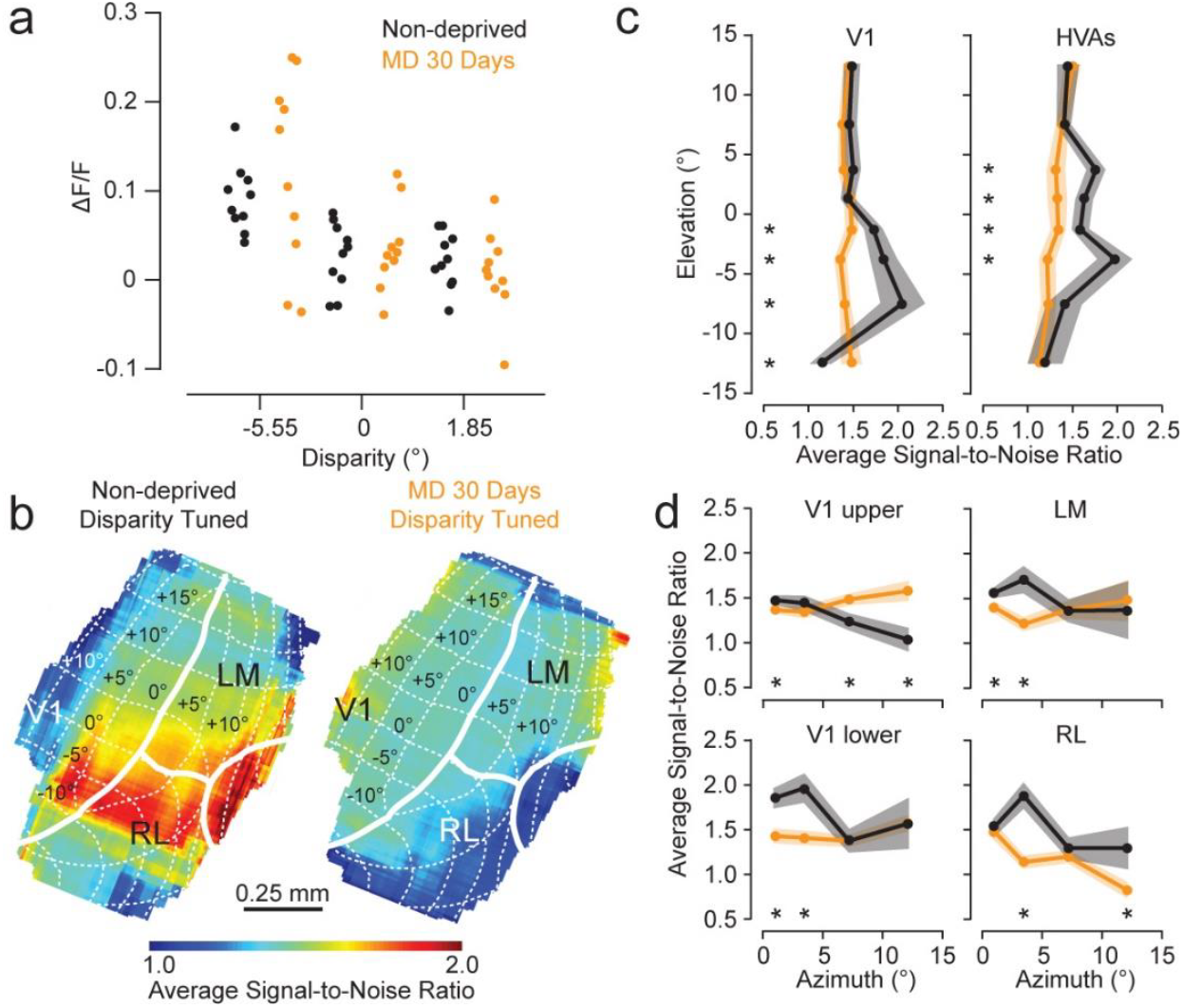
Monocular deprivation severely reduced signal-to-noise ratio of responses to disparity. **a**, Single-trial responses to three different disparities for two example neurons (n = 10). **b**, Each point in the map represents the average signal-to-noise ratio (SNR, mean/standard deviation) of significant disparity-tuned neurons within a ±0.25 mm window for either non-deprived (left) or MD (right) mice. Each pixel in the maps included at least 10 significant disparity-tuned neurons in both non-deprived and 30-day MD mice. **c**, Mean SNR for non-deprived mice (black) and 30-day MD mice (orange) with respect to elevation for neurons in V1 (left, bootstrap ANOVA, p < 0.0001 and p = 0.26, respectively) and HVAs (right, p = 0.01 and 0.03, respectively). **d**, Mean SNR for non-deprived mice (black) and 30-day MD mice (green) with respect to azimuth for neurons in upper V1 (upper left, bootstrap ANOVA, p = 0.001 and 0.07, respectively), lower V1 (lower left, p = 0.002 and 0.07, respectively), LM (upper right, p = 0.08 and 0.02, respectively), and RL (lower right, p = 0.001 and p < 0.0001, respectively). Significant differences (bootstrap test, p < 0.05) between non-deprived and 30-day MD data points are noted with an asterisk. Shaded regions are standard error of the mean.

To demonstrate how this degraded organization explains impaired binocular depth performance in the PDCT, we discriminated disparities that correspond to depths encountered in the task using single-trial neuronal responses. Previously, we calculated what the disparities would be for the depths of near and far surfaces in the PDCT based on the position and behavior of the mice during the task^29^ (Figure 7a). Using all neurons that would detect the near surface (preferred disparity ranging from -7.37° to -3.70°), we computed d’ comparing the mean and trial-to-trial standard deviation of responses to -5.55° (closest disparity to a depth of 2.5 cm) and either 0° (closest disparity to a depth of 15.2 cm) or 1.85° (closest disparity to a depth of 30.5 cm). We found that similar to the behavior, individual neurons from non-deprived mice significantly discriminated -5.55° from 0° better than mice with 30 days of MD (Figure 7b, left). For the larger difference in disparity, d’ was larger for both non-deprived and 30-day MD mice and the difference between non-deprived and 30-day MD mice was smaller, but still significant (Figure 7b, right).

**Figure 7.**
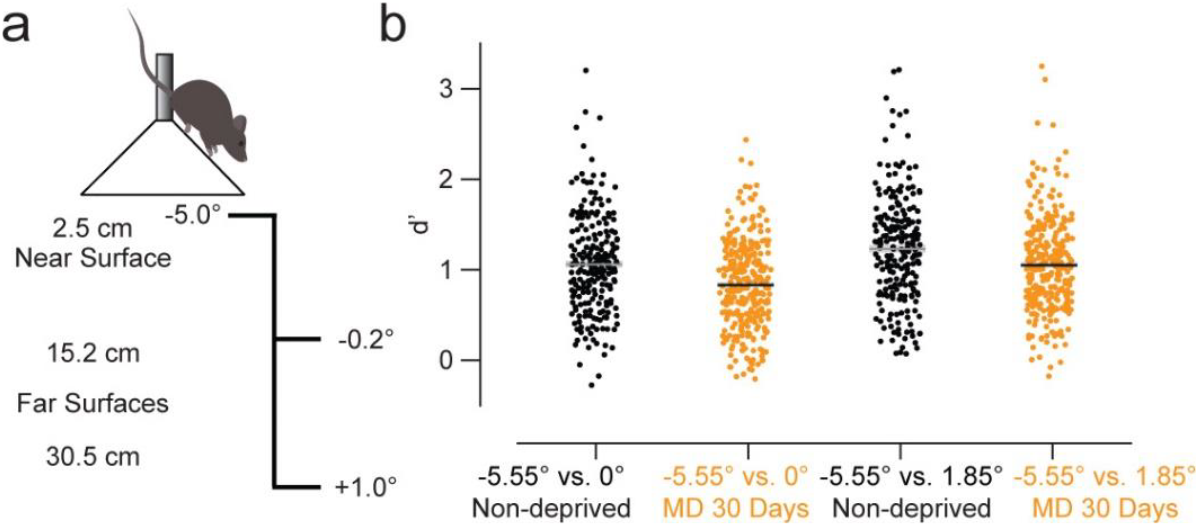
Neuronal responses can predict deficits in pole descent cliff task behavior. **a**, The near surface in the pole descent cliff task corresponds to a disparity of -5° and the two far surfaces correspond to disparities of -0.2° and +1°, respectively^29^. **b**, Trial-to-trial disparity discrimination (d’) based on single neuron responses tuned to the near surface was reduced in 30-day MD mice compared to non-deprived mice (bootstrap test, n = 219 and n = 307, p < 0.0001 and p = 0.0004 for fine and coarse disparity differences, respectively). Shaded regions and error bars are standard error of the mean with respect to neurons.

The results of the PDCT in Figure 1 are based on 15 trials and each point represents a single mouse. There is considerable variance in task performance on the PDCT and thus we measured the performance of ∼25 mice per group. We measured neuronal responses in only a small subset of these mice because not all mice expressed GCaMP6s, so we did not evaluate direct relationships between the PDCT and neuronal d’ data.

## Discussion

Monocular deprivation spanning the critical period results in an enduring deficit in binocular depth discrimination (Figure 1b). Previously we demonstrated that depth discrimination with the PDCT could be explained by the disparity tuning in the visual cortex^29^. One previous study measured weaker disparity selectivity in mice immediately following 8 days of MD^28^. Thus, the first hypothesis we formulated was that disparity tuning would be weaker in mice receiving 30 days of MD. However, we did not detect any reduced disparity selectivity in 30-day MD mice. Mice receiving 30 days of MD followed by several weeks of binocular vision had similar visual responsiveness (Figure 2), similar percentages of significant disparity-tuned neurons (Figure 2), a similar representation of a broad range of disparities (Figure 4), and similar disparity selectivity (Figure 5) compared to non-deprived mice. We conclude that although 30-day MD mice possess the neuronal response characteristics sufficient to discriminate binocular depth, they exhibit a behavioral deficit. We argue that the reduced ability to discriminate depth is a consequence of the disrupted mapping of disparity-tuned neurons in the central portion of their visual field predominantly in HVAs.

The central portion of the visual field has been proposed to represent an enhanced region in the mouse with tighter retinotopic organization in the visual cortex^36^. Mice shift their gaze with head and eye movements to put objects of interest within this portion of the visual field^37-39^. We found that mice also have a larger percentage of disparity-tuned neurons (Figure 3), preferentially represent near surfaces within this region (Figure 4), and have more reliable trial-to-trial responses to disparity (Figure 6). Mice receiving 30 days of MD displayed a broader spatial representation of the ratio of disparity-tuned neurons and changes in preferred disparity, as well as lower trial-to-trial reliability compared to non-deprived mice. Previous studies in the cat and ferret have shown that altered visual experience can severely disrupt the formation of feature maps in V1^34,40^ and trial-to-trial reliability is enhanced over the critical period of development^35^. This broader spatial representation and lower trial-to-trial reliability might explain why 30-day MD mice perform worse in the PDCT, which was supported by the reduced discriminability of near from far surfaces in individual near-tuned neurons (Figure 7). We propose that the function of the disparity maps is to cluster neurons with similar preferred disparity together to improve the reliability of responses and support binocular depth discrimination. Our analysis does not rule out additional contributions to reduced PDCT performance. Since 30 days of MD also results in reduced spatial acuity^18^, it would be interesting to examine direct relationships between spatial and disparity properties in future experiments.

The map in Figure 4 is based on average preferred disparity. We find neurons with a broad range of preferred disparities at each point in the map, but the average preferred disparity varies systematically with respect to elevation. Therefore, it is not as uniform as maps based on preferred orientation measured in the visual cortex of mammals such as ferrets, cats, and non-human primates^1,2,34,40^. However, the disparity map in mice is very similar to how preferred disparity is distributed in V1 of non-human primates suggesting that our results could be applicable to many other species^27^.

We also observed that the disparity-dependent mapping was stronger in HVAs in non-deprived mice compared to mice after 30 days of MD. One explanation of higher SNR could be that some neurons in non-deprived mice are more tolerant to trial-to-trial variations in binocular alignment. Humans depend on relative disparity cues for fine binocular depth discrimination because it does not depend on precise binocular alignment^41-43^. V2 in the non-human primate responds more selectively to relative disparity or disparity edges compared to V1, which responds primarily to absolute disparity^44-46^. The HVAs are secondary visual areas in the mouse that could be analogous to V2 in the non-human primate. Recent work has proposed that HVAs are important for simple and complex behaviors^47,48^. Mice have considerable variation in their binocular alignment so relative disparity tuning would improve binocular depth discrimintation (Extended Data Figure 3). Previously, we determined that mice target the edges between platforms when performing the PDCT^29^. The stimuli used in these experiments included both absolute and relative disparity information and additional experiments are needed to determine whether relative disparity selectivity contributes to this reduced variability observed in HVAs.

Overall, we find that disrupting visual experience during the critical period causes severe degradation of the retinotopic and hierarchical organization of disparity tuning properties in V1 and HVAs. We demonstrate how and why both the behavioral performance and neuronal discrimination are reduced in mice lacking proper mapping. Thus, we conclude that such mapping plays an important role in binocular depth perception.

## Acknowledgments

We thank Céleste-Élise Stephany for performing the classic visual task on non-deprived and 30-day MD mice. We are grateful to Ian Nauhaus for providing the software and equipment needed for retinotopic mapping and Nicholas Priebe for sharing the equipment and resources necessary to carry out the two-photon experiments. We thank Mandi Severson and Ronan O’Shea for providing technical assistance and Boris Zemelman and Stefanie Esmond for providing genotyping. Funded by NIH R01EY034092 (AWM & JMS).

## Materials and Methods

### Preparation of animals

All procedures were approved by The University of Texas at Austin Institutional Animal Care and Use Committee (IACUC) and the University of Louisville School of Medicine IACUC, the University of Arizona College of Medicine - Phoenix IACUC, and are in accordance with the National Institutes of Health Guide for the Care and Use of Laboratory Animals. For two-photon microscopy, we used tetO-GCaMP6s mice (Jackson Labs, # 024742) crossed with CamKII-tTA mice (Jackson Labs, # 007004) to express the GCaMP6s calcium indicator in excitatory neurons in 26 mice^31^. Ten of these mice had normal binocular vision during the critical period (non-deprived), including 7 males and 3 females. The other 16 mice (11 males and 5 females) had their right eye sutured closed at postnatal day (P) 22 under isoflurane anesthesia (1–3%) and removed under anesthesia following 30 days of monocular deprivation (MD) (P52) as described in detail previously^18,30^. For these mice, as described previously^32^, a titanium bar was secured to the skull using dental acrylic and a 3- or 4-mm craniotomy was made over the binocular region of the left hemisphere of V1 and a glass window was secured in place with cyanoacrylate under isoflurane anesthesia (1–3%). For the pole descent cliff task (PDCT), we used 97 total mice with 51 that underwent the same 30-day monocular deprivation procedure used for two-photon microscopy. These mice were habituated to handling for two consecutive days and had their whiskers clipped a day before performing the task. All mice were 3-7 months in age when imaged or tested on the pole descent visual cliff task. For the classic visual cliff tasks, we used 35 total mice with 11 that underwent the same eye suture procedure as described above just before being tested, and another 13 mice that underwent the same 30-day monocular deprivation procedure as described above.

### Pole descent cliff task

Following the methods outlined in [29], we tested non-deprived and 30-day MD mice on the PDCT. In the PDCT, the mouse descends a vertical pole to the glass divided into four quadrants (Figure 1A). One quadrant is closer, with a 2.5 cm surface below the glass, whereas the three other quadrants have surfaces with deeper depths. A 9.5 cm diameter cone rests at the bottom of the pole on thin metal rails

1.5 cm above a plate of glass and 2.5 cm above the nearest platform. These rails obscure the edges between the quadrant presenting the nearest platform and the remaining three quadrants. All interior surfaces are covered with a black and white 2.5 cm checkerboard pattern. We place mice on the pole and then measure the fraction of 15 trials that each mouse exits the cone onto the nearest (2.5 cm) surface as the fractional success (Figure 1B). For a subset of these mice, we also measured the time it took to descend the pole and choose a surface (Extended Data Figure 1).

### Classic visual cliff task

The classical visual cliff test has traditionally been used for testing depth perception in multiple species^49^. Following the methods outlined in^50^, we tested mice on the visual cliff task. In the visual cliff task, a mouse is placed on a square aluminum rail 3.8 cm above a plate of glass (Extended Data Figure 2). Immediately under the glass on one side is a surface covered in 2.5 cm black and white checkered cloth. On the other side, this surface is positioned 61cm below the glass plate. Mice were tested for ten interleaved trials. The position of the closest surface was determined at random for each trial number (1-10). Mice were placed on the glass and scored as to which side of the rail they exited. The glass and rail were cleaned after each trial with a glass cleaner solution.

### Retinotopic mapping

Fluorescence was measured in the entire imaging window using the same intrinsic optical imaging and stimulation setup described previously^33,51,52^. For visual stimuli, we used a contrasting checkered bar^33^ presented for twelve trials, alternating between the four cardinal directions at random projected onto a 64 × 46-cm screen using a LC4500-UV projector (Keynotes Photonics). The screen was placed 10 cm away and centered on the right eye at a 30-degree angle. While imaging, the mice were head-fixed with a custom stand and lightly anesthetized using 0.5% isoflurane. Silicone oil was applied to the eyes. We used a pco.panda 4.2 sCOMAS camera (Excelitas) fitted with a 25 mm C VIS-NIR fixed focal length lens (Edmond Optics) and a 525 nm fluorescence filter (Thor Labs). Clay was used to block out external light for the region between the imaging window and the lens. To illuminate the cortex, we used an X-Cite 110 LED illuminator (Excelitas) with a 460 nm filter (Thor Labs). The V1 and RL/LM border was identified in individual mice based on the retinotopic reversal in the azimuth dimension and the RL and LM border was identified by a retinotopic reversal in the elevation dimension in AL (anterolateral area). Once all two-photon data was aligned based on these two borders, we used borders and retinotopic contours that were previously averaged over several mice for identifying retinotopic location and area identification^33^.

### Two-photon microscopy

Changes in fluorescence from the calcium indicator GCaMP6s expressed in excitatory neurons in CamKII-tTA/tetO-GCaMP6s mice^31^ were measured using similar methods as those described in detail previously^32^. For stimuli, we presented 6-s dynamic random dot stereograms (DRDS) (6 Hz, 7.4° dots, 200 black and 200 white dots) interleaved with 6-s mean gray screen with Psychtoolbox^53^ using an Optoma HD27 projector with a refresh rate of 120 Hz to rear-project images onto an RP3D polarization-preserving screen (Severtson Screens) placed 22 cm in front of the mouse. Stereoscopic depth was generated by presenting shifted dots to each eye every other frame using a circular polarization alternator (DepthQ/Lightspeed Design) synchronized with the projector refresh rate and passive circular polarization filters placed in front of the mouse eyes. The DRDS was 70° wide and 100° high and binocular disparity was varied at 0°, ±1.85°, ±5.55°, and ±9.20° in the vertical central 65% portion and remained at 0° on each side (10 different DRDS for each disparity). For imaging, we used a custom-built two-photon resonant mirror scanning microscope with a mode-locked 920-nm Chameleon Ultra Ti:Sapphire laser (Coherent Technologies)^23^. Clay was used to block out external light. Green light focused by a 16× water objective (0.8 numerical aperture; Nikon) through Aquasonic Clear ultrasound gel (Parker Laboratories) was collected from a 0.4 to 0.55 mm-square region with photomultiplier tubes. Images (256 × 455 pixels) were obtained with custom software at a frame rate of 30.9 Hz (Labview; National Instruments). The objective was rotated normal to the cortical surface and focal planes ranged from 130 to 200 μm below the cortical surface.

During imaging, we also tracked the pupils of both eyes as described previously^32,54,55^. Two infrared cameras (30 frames per second) were mounted in front of each eye perpendicular to the orbital axis while the mouse was head-fixed and running on a floating trackball (Dombeck et al. 2007). Pupil position for each eye was tracked with DeepLabCut^56^. Position was calibrated to degrees of visual angle based on an artificial eyeball of similar size with known rotations. There was no significant different in variation in trial-to-trial eye position for MD 30-day mice compared to non-deprived mice (Extended Data Figure 3a-b) and there was no significant difference in binocular alignment between non-deprived and 30-day MD mice (Extended Data Figure 3c-d). As previously found, eye position did not depend on disparity^32^ (Extended Data Figure 3e-h).

### Two-photon data analysis

Images were analyzed with custom Matlab software and were first bandpass filtered over time using a moving average of 8 frames and subtracting a median filter of 1000 frames. Neurons were identified by hand from images and video based on changes in brightness, size, and shape. Masks were drawn around neurons and any overlap between masks was discarded. The neuropil was defined as all pixels outside of all masks. A neuropil annulus for each mask was defined as an annulus 4 pixels wide and 3 pixels from the mask along the 455-pixel dimension and 2 pixels wide and 2 pixels away from the mask along the 256-pixel dimension (any overlap with other masks discarded). We then computed the mean brightness of all pixels within each mask, the neuropil, and neuropil annuli. Then, we subtracted the average intensity calculated 4 seconds before all stimuli from these intensities (Δ*F*). Then we divide those differences by the median intensity (*F*_0_) over the entire experiment to get a Δ*F*/*F*_0_. For masks, we then additionally subtract the neuropil annulus value for each mask for a final neuropil corrected Δ*F*/ *F*_0_.

Visually-responsive neurons were defined as those neurons with a significantly greater average responses for the 6 s during the stimulus compared to 4 s before the stimulus (sign test, p > 0.05, n = 70 trials). Disparity-tuned neurons were defined as those neurons with a significantly different average response with respect to disparity (Kruskal-Wallis test, p < 0.05, n = 10 trials, 7 disparities). The disparity selectivity index (DSI) was defined as the resultant vector calculation^57^:

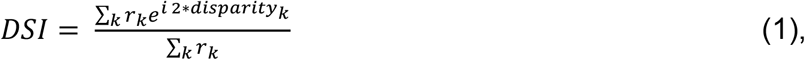

where *r*_k_ is the response to each disparity. Since this is a circular calculation, we assumed that the disparities we tested wrap around at the extremes. If all responses are equal, DSI = 0 and if the neuron only responds to a single disparity, DSI = 1. Preferred disparity was defined as the disparity that produced the maximum response based on a Gabor function fit to the average responses to the 7 disparities:

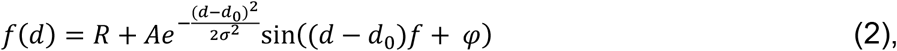

where *d* is disparity. We only used the Gabor function to determine preferred disparity if the function explained at least 50% of disparity-dependent variance and after visual inspection that the value was consistent with the data. Otherwise, whichever of the 7 disparities produced the largest average response was assigned as the preferred disparity for that neuron. On average, 84% of the Gabor fits were able to explain at least 50% of the variance.

To generate cortical maps of ratio of preferred disparity and preferred disparity, all neurons were assigned a location in the window based on the center of the mask within the two-photon region-of-interest (ROI). Then, the center of the ROI was assigned a global location with respect to the V1/RL/LM border. If necessary, locations were also translated for orientation to account for any rotation to align with the V1 border. Once all locations were aggregated for either non-deprived or 30-day MD mice, all points were translated with respect to a single rotation so that the V1 border was approximately vertical and the RL/LM border was approximately horizontal. Then we computed either the ratio of neurons, average preferred disparity, DSI, or SNR within a sliding ±0.25 mm window along these axes every 0.01 mm. The final map only included pixels where there were at least 100 neurons or 10 disparity-tuned neurons within the window of both non-deprived and 30-day MD mice. The final maps were then rotated back to align with the average V1 border and to align with a medial-to-lateral (left-to-right) and posterior-to-anterior (top-to-bottom) orientation^33^.

To plot the ratio of neurons, average preferred disparity, DSI, or SNR with respect to elevation or azimuth, we converted the cortical location (in mm) of each neuron into a retinotopic location (degrees of visual angle). For azimuth, the V1/HVA border was defined as 0° and used to assign retinotopic coordinates as V1 or the HVA areas of RL or LM. For elevation, 0° was defined by the V1/RL/LM border. Anything above the elevation of this point was defined in positive values and as upper V1 or LM and anything below the elevation of this point was defined in negative values and as lower V1 or RL. We used a single complete retinotopic map based on the average elevation and azimuth maps across mice using the procedure described above. We used an average map to reduce retinotopic map noise as much as possible in this translation procedure. Retinotopic maps were measured using a different setup than what was used during two-photon microscopy and optical imaging produces a signal with relatively low signal-to-noise compared to two-photon microscopy. There were some distortions between the vasculature in the images produced by each setup due to optical equipment and angular alignment differences so we prioritized aligning two-photon windows with respect to vasculature closest to the V1/RL/LM border for each mouse. Despite this individual variability and potential uncertainty, variations that we observed in the final cortical maps of the ratio of neurons, average preferred disparity, DSI, or SNR measured with two-photon microscopy appeared to align very well with the contours of average retinotopic maps (e.g., Figure 3b, left).

For single-trial discrimination, we used neurons with preferred disparities ranging from -7.37° to -3.70° and d’ was calculated for each neuron, individually, with *µ* and *σ* calculated across the 10 repeats for each disparity:

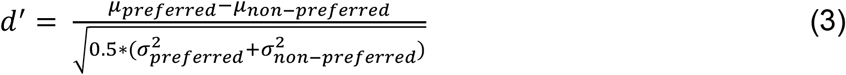

We then averaged d’ across the population. Since the disparities we used for imaging do not match the depths of the pole descent cliff task exactly, we discriminated the closest disparities that were shown to the mice (preferred = -5.5°; non-preferred = 0° or 1.85°).

### Statistics

All statistical tests were done in Matlab and the details of the tests are included in the text for figure legends. We did not use a specific analysis to determine population size, but the number of animals, trials, and neurons are similar or larger than those used in previous studies^29,32^. Data collection and analysis were not performed blinded to subject conditions. No assumptions were made about data distributions and statistical tests of individual neurons were non-parametric: Kruskal-Wallis for disparity tuning and sign test for visual responsiveness. For all other tests, we used bootstrap analysis of the mean (*µ*)^58^. For a bootstrap test between two sets of data, we resampled each set of data 20,000 times, allowing repeats, to produce surrogate datasets of the same size. Sorted estimates from these datasets (percentiles) were then used to determine a confidence interval and p values relative to a null hypothesis (*µ*_1_ = *µ*_2_). If the entire surrogate datasets were above or below the null hypothesis, we described the result as p < 0.0001. For a bootstrap multiple comparison test between multiple sets of data (bootstrap ANOVA), we resampled each set of data 20,000 times, allowing repeats, to produce surrogate datasets of the same size and resampled the combined set of data 20,000 times, allowing repeats to produce surrogate datasets of the combined data. Sorted estimates from each of *n* datasets and the combined (*all*) dataset (percentiles) were then used to determine p values relative to a null hypothesis (*µ*_1_ = *µ*_all,_ *µ*_2_ = *µ*_all, …,_ *µ*_n_ = *µ*_all_). The p value was the lowest result of comparisons between the *n* datasets and the combined dataset. Again, if the entire surrogate datasets were above or below the null hypothesis, we described the result as p < 0.0001. All data for all figures is available at figshare: https://figshare.com/projects/Degraded_mapping_of_disparity_tuning_in_visual_cortex_explains_defecits_in_binocular_depth_perception/245687

---

**Extended Data Figure 1.**
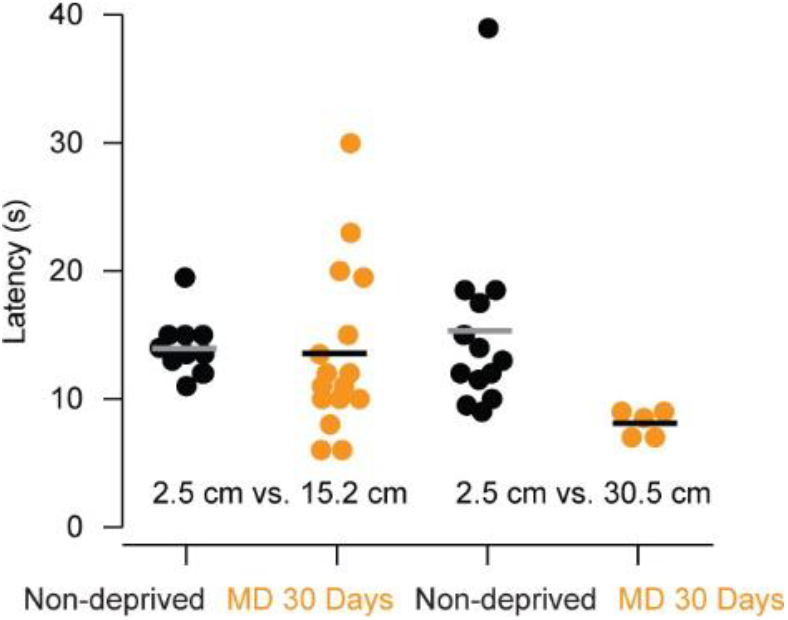
30-day MD mice do not take longer to descend the pole and choose a surface compared to non-deprived mice. These mice (each data point) are a subset of the mice included in Figure 1b.

**Extended Data Figure 2.**
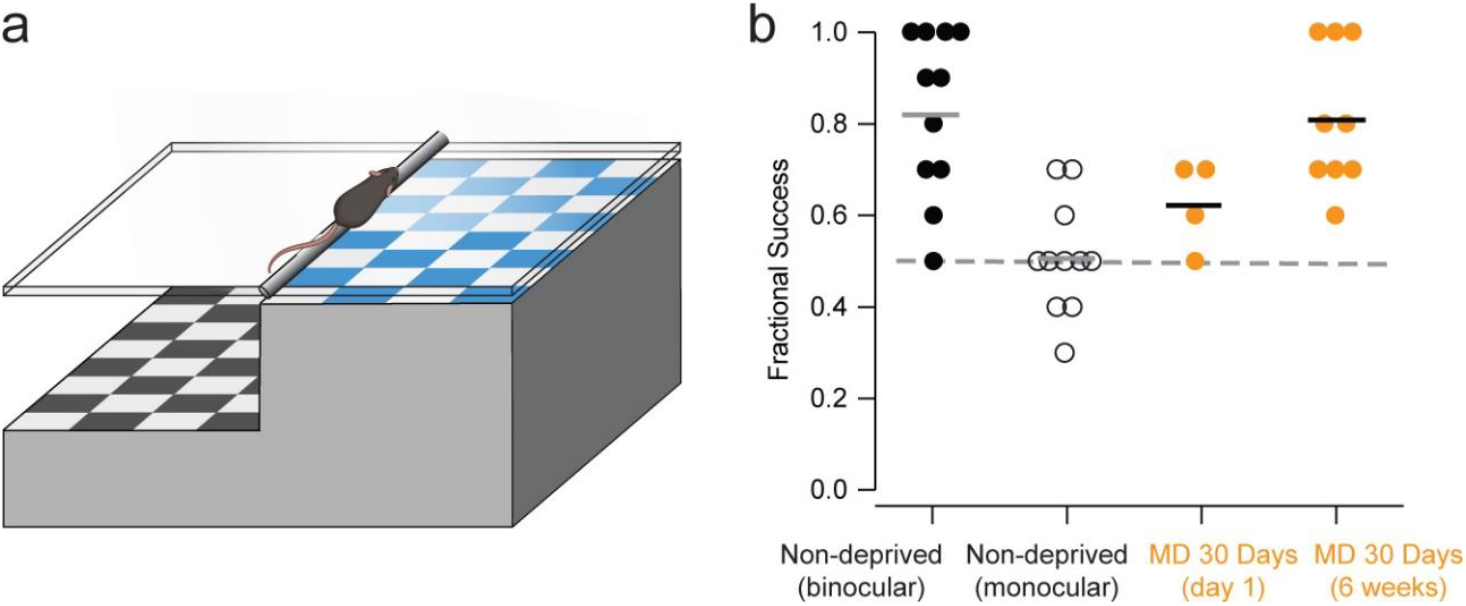
Long-term monocular deprivation does not result in persistent impaired binocular depth discrimination in the classical visual cliff task. **a**, The classic visual cliff task was used to measure binocular depth discrimination in non-deprived and 30-day MD mice. **b**, The fraction of trials that each mouse chose the near surface (2.5 cm) versus the far surface (61 cm) when placed on glass over the border of a visual cliff apparatus (Wilcoxon signed rank test, p = 0.002, N = 11 and 11 mice). Non-deprived monocular mice that had one eye sutured closed (well after the critical period) and were tested immediately after recovering from the procedure did not choose the near surface significantly beyond chance (dashed line, Wilcoxon signed rank test, p = 1.0, N = 11 mice). Four mice with 30 days of MD over the entire critical period appeared to have a deficit (although not significant with such a small sample) one day after reopening the closed eye (Wilcoxon rank sum test, p = 0.08). There was no significant difference compared to deprived mice for a larger number of 30-day MD mice after 6 weeks of normal binocular vision after reopening the closed eye (Wilcoxon rank sum test, p = 0.84, N = 9 mice).

**Extended Data Figure 3.**
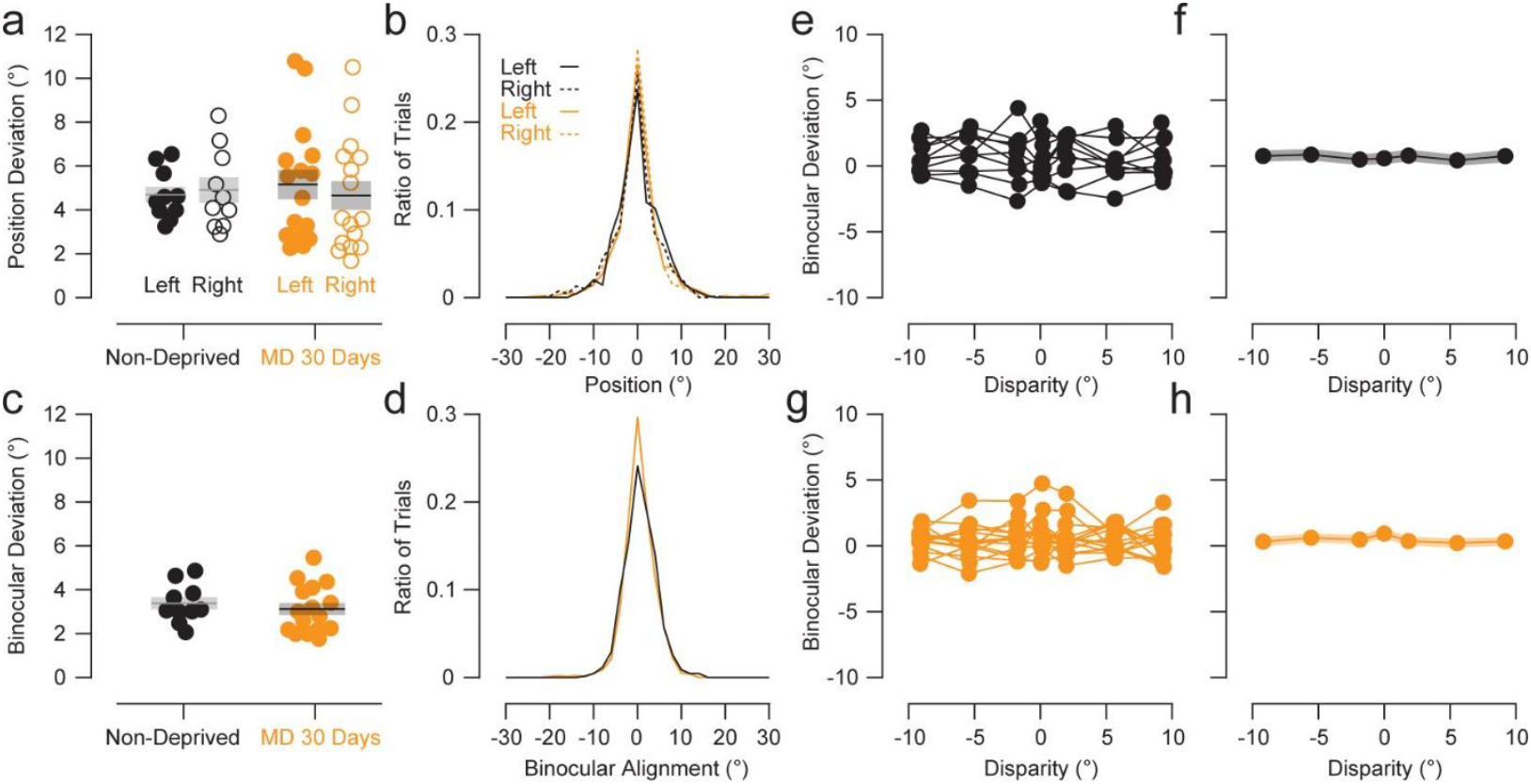
Binocular eye position variation during dynamic random dot stereogram stimulation. **a**, Standard deviation of eye position across trials (n = 70) for each eye in each mouse. Variation was not significantly different between non-deprived and 30-day MD mice (bootstrap test, p = 0.48 for left eye and p = 0.67 for right eye, N = 10 and 16 mice). **b**, Eye position data for all mice combined into a single distribution. **c**, Standard deviation of binocular alignment (left – right eye) across trials (n = 70) in each mouse. Variation was not significantly different between non-deprived and 30-day MD mice (bootstrap test, p = 0.33, N = 10 and 16 mice). **d**, Binocular alignment data for all mice combined into a single distribution. **e**, Binocular alignment separated by which disparity was shown (n = 7 disparities, 10 trials). **f**, The standard deviation of binocular alignment did not depend on disparity for alignment data combined for all non-deprived mice (bootstrap ANOVA, p = 0.51). **g**, The same values as **e** for 30-day MD mice. **h**, The standard deviation of binocular alignment did not depend on disparity for alignment data combined for all 30-day MD mice (bootstrap ANOVA, p = 0.13).

